# The glue produced by *Drosophila melanogaster* for pupa adhesion is universal

**DOI:** 10.1101/2019.12.19.882761

**Authors:** Flora Borne, Alexander Kovalev, Stanislav Gorb, Virginie Courtier-Orgogozo

## Abstract

Insects produce a variety of adhesives for diverse functions such as locomotion, mating, egg or pupal anchorage to substrates. Although they are important for the biology of organisms and potentially represent a great resource for developing new materials, insect adhesives have been little studied so far. Here, we examined the adhesive properties of the larval glue of *D. melanogaster*. This glue is made of glycosylated proteins and allows the animal to adhere to a substrate during metamorphosis. We designed an adhesion test to measure the pull-off force required to detach a pupa from a substrate and to evaluate the contact area covered by the glue. We found that the pupa adheres with similar forces to a variety of substrates (with distinct roughness, hydrophilic and charge properties). We obtained an average pull-off force of 217 mN, corresponding to 15 500 times the weight of a pupa and adhesion strength of 137-244 kPa. Surprisingly, the pull-off forces did not depend on the contact area. Our study paves the way for a genetic dissection of the components of *Drosophila melanogaster* glue that confer its particular adhesive properties.

**SUMMARY STATEMENT:** We designed an adhesion test to measure the pull-off force required to detach drosophila pupae and found that Drosophila glue adheres similarly to various substrates of different chemical properties.

## INTRODUCTION

Natural adhesives have a very important significance for the biology of organisms and are a great material for innovation of biologically-inspired technical adhesives. The most studied bioadhesives are from marine organisms, mussels and barnacles (Power et al., 2010), and they are now used in a variety of biomimetic applications such as surgical sealants to repair tissues or synthetic polymer coatings (Lee et al., 2011). Bioadhesives in insects are less studied although they are extremely diverse (Gorb, 2001; Graham, 2008; Li et al., 2008; Betz, 2010). Some insects produce glue to secure their eggs or cocoon (Betz, 2010). A few studies have been carried out on egg glue composition and particularly on the glue adhesive strength in different insect species. For example, in *Opodiphthera* moths, females secrete a viscous fluid from their accessory reproductive gland that sticks their eggs to each other and to the substratum. This hydrogel-type glue is highly elastic and mainly made of proteins. The shear strength reaches 1-2 MPa on wood, which could potentially be enough for some industrial applications (Li et al., 2008). Furthermore, these eggs and eggs from other species such as *Crioceris asparagi* have the ability to attach to plants covered with crystalline epicuticular waxes known to be non-adhesive (Voigt and Gorb, 2010). Similar properties have been revealed in the egg glue of codling moth (Al Bitar et al., 2012; 2014). Thus, investigating the glue of various insect species appears to be a great strategy to find novel universal and substrate-specific adhesives. Surprisingly, although *Drosophila melanogaster* is a model organism that is extensively used in laboratories, little is known about pupal adhesion in this species.

*Drosophila melanogaster* larvae secrete a glue from their salivary glands right before pupariation (Fraenkel and Brookes, 1953). After expectoration, following larval peristaltic movements, the secreted liquid spreads between the body and the substrate and dries within a few minutes (Beňová-Liszeková et al., 2019). This glue allows the animal to stay firmly attached to an external surface for several days, until the adult fly emerges from the pupal case, while the pupal case remains attached to the substrate. The pupa of other Brachycera fly species is also often attached to a substrate during metamorphosis (Fraenkel and Brookes, 1953).

In the wild, *Drosophila* pupae have been reported to adhere to a great variety of substrates from fruits to beer bottles (Vouidibio et al., 1989) and in diverse environments. Particularly, pupae have been observed on dry parts of various fruits, some species have also been found fixed to wood, wet rotten parts of fruits, deep in the soil and to one another (Grossfield, 1978; Sokolowski, 1985; Vandal et al., 2008; Beltramí et al., 2010; Castillo et al., 2014). Pupal attachment might be important for not being taken away by predators, to resist environmental conditions (wind or rain) or to help adult flies emerge from the pupal case after metamorphosis (Da Lage et al., 2019).

In *D. melanogaster* (Korge, 1975; 1977) and other *Drosophila* species (*D. virilis* (Kress, 1982), *D. natusa* (Ramesh and Kalisch, 1988), *D. gibberosa* (Shirk et al., 1988)), the glue is composed of a small number of proteins called salivary gland secreted (Sgs) proteins. These proteins present repeated motifs and glycosylations, which are commonly found in adhesive proteins (Graham, 2008; Betz, 2010). Some of the *Drosophila* glue proteins are rich in cysteine, like many marine adhesive proteins and this property could be important to maintain a secondary structure by forming disulfide bonds (Graham, 2008). Other *Drosophila* glue proteins are rich in serine and threonine and highly O-glycosylated, suggesting that they may interact with water to hydrate or dehydrate the glue (Farkaš, 2016). The *Sgs* genes have evolved rapidly between species and within species (Korge, 1975; 1977; Beckendorf and Kafatos, 1976; Da Lage et al., 2019).

Here, we investigated the adhesive properties of the glue of *D. melanogaster* on different substrates. First, we analyzed the contact region between the pupa and the surface to which it is attached. Then, we designed a pull-off force measurement set up. We assessed the force required to detach the pupa from different substrates, with different hydrophilic and charge properties or with different roughnesses, and analyzed the way rupture occurs.

## METHODS

### Flies

*D. melanogaster* flies were cultured in plastic vials on standard medium (1L: 62.5 g yeast, 62.5 g cornmeal, 10.0 g agar-agar, 20.0 g glucose monohydrate, 30.0 g molasse, 30.0 g sugar beet syrup, 10.0 ml propionic acid (10%), nipagin (10%)) at room temperature. The stock Canton S (gift from Roger Karess) was used for force measurement assays and SEM microscopy. The stock *w[*]; P{w[+m*]=Sgs3-GFP}2* (FBti0016953; Bloomington Drosophila stock #5884; Biyasheva et al., 2001) was used for confocal microscopy.

### Larva preparation

Third instar wandering larvae were washed in PBS to remove traces of food and microorganisms from their surface, put in empty Petri dishes with a paintbrush and kept in high humidity atmosphere at room temperature in a closed plastic box (15×10×5 cm) containing wet cotton. When larvae stopped moving, they were transferred on the substrate of interest with soft forceps, kept at high humidity as mentioned above and let to pupariate. Five to 24 h after transfer to the substrate of interest, the substrate with pupae attached to it was put for 1 h at room humidity to allow the glue to dry completely and then the pupae were used for adhesion assays. Pupae not used for assays were weighed individually using Mettler Toledo AG203 (DeltaRange®, Gießen, Germany). Temperature, humidity and atmospheric pressure were monitored daily.

### Preparation of glass and Teflon substrates

Two types of non-coated microscopic glass slides were used to measure adhesion on glass: Menzel Superfrost microscope glass slide from ThermoScientific™ (#AGAB000080) and microscopic glass slide from Roth (#0656.1). Atmospheric plasma-treated glass slides were prepared using a PlasmaBeam (Diener electronic GmbH, Ebhausen, Germany) on Roth glass slides for 1 min. To prepare Poly-L-Lysine-Polyethyl glycol-coated (PLL-PEG-coated) glass slides, Roth glass slides were cleaned with a plasma cleaner then coated with 0.1 mg·ml^−1^ non-biofunctionalized PEG sidechains (methoxy-terminated) (SuSos). Poly-L-Lysine-coated (PLL-coated) glass slides from Thermo Scientific™ (J2800AMNZ) and pieces of Teflon (Polytetrafluorethylen, Kelux, Geldern, Germany) were also tested.

### Preparation of Spurr epoxy resin substrates

First, a silicone cast was made by pouring polyvinylsiloxane (light body Affinis, Coltène/Whaledent GmbH + Co. KG, Langenau, Germany) over a cleaned Roth glass slide (to create the smooth resin, Ra=80 nm (Salerno et al., 2014)) or over glass slides covered by polished papers with different grain sizes: 1 μm and 9 μm (Ra=0.54 and 2.47 μm measured over 1400×1050 μm area using NewView 6k (Zygo, Middlefield, CT, USA) white light interferometer; both with FiberMet Abrasive disks, Buehler) (Salerno et al., 2014), P1000 and P80 (Ra=3.94 and 40.40 μm measured over the same area using VR 3100 (Keyence, Neu-Isenburg, Germany) 3D profilometer; polishing papers, Bahaus). After a couple of minutes, the slide and the polished paper were removed and the edges of the polyvinylsiloxane cast were made higher. Then, Spurr epoxy resin (Spurr, 1969) was poured into the polyvinylsiloxane cast and polymerized at 65 °C overnight in Memmert U 15 oven (Schwabach, Germany). Resin was then allowed to cool down for a couple of hours before unmolding. Each resin was used for several assays.

### Contact angle measurements

Water contact angles on different substrate surfaces were measured using a contact angle measurement device OCA20 (DataPhysics Instruments, Filderstadt, Germany). A 2-μl water droplet was deposited on a substrate, then an image of the droplet was taken after five seconds, and contact angles were determined from the fit of the droplets shape with a sphere using SCA 202 software (DataPhysics Instruments, Filderstadt, Germany). All the contact angle measurements were done before the pull-off force assays except for PLL-PEG-coated glass.

### Adhesion force measurement

To measure pupal adhesion, a force transducer (100 g capacity, FORT100, World Precision Instruments, Sarasota, FL, USA) mounted on a motorized 3D micromanipulator (DC3001R, World Precision Instruments, Sarasota, FL, USA) was used (Fig. 1). The substrate was horizontally clamped to a lab boy. A piece of aluminum platelet of 1.0 x 2.0 x 0.2 mm attached to the force transducer by a metal wire (0.1 mm) was covered by a piece of double-sided adhesive tape (tesa, extra strong, #05681-00018). A new piece of tape was glued for each measurement. Only pupae attached on their ventral side and which did not contact other pupae were used for measurements. The tape on aluminum platelet was brought into contact with the dorsal part of the pupa by adjusting manually the height of the lab boy. The force transducer was then moved upwards with a velocity of 200 μm/s until detachment of the pupa. The signal from the force transducer was amplified using Transbridge TBM4M and digitized using Lab-Trax-4 data acquisition hardware (WPI, Sarasota, FL, USA). Three phases could be distinguished on force curves recorded using LabScribe v3 (iWorks, Dover, NH, USA) (Fig. 2A). During the initial phase, the pupa was not attached to the tape and force with no pupa weight was measured. The pulling phase started when the pupa came into contact with the tape. During this phase, the pupa was stretched until it detaches from the substrate (maximal force). Finally, during the resting phase, the force went down to a basal value including pupal weight. Overall, the adhesion strength we measured correponds to the weakest interface: adhesion of the glue to the substrate, cohesion of the glue, adhesion of the glue to the pupal case, cohesion of the pupal case (when the cuticle broke) or the interface between the tape and the pupal case (when the tape detached from the pupa). The set up was calibrated before the first assay with a standard weight. Pull-off force corresponding to the force at pupa detachment was calculated as the difference between the maximal pulling force and the mean force at the initial phase (offset). The force corresponding to the adjustment of the lab boy until pupa glued to the tape between the initial phase and the pulling phase was not recorded.

**Figure 1.**
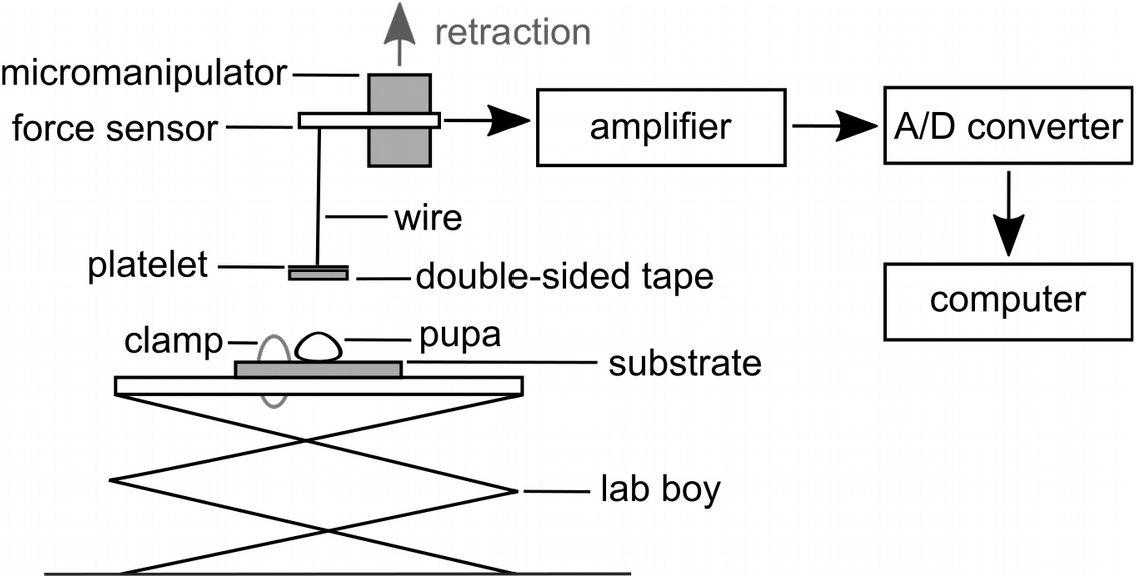
Experimental set-up for measuring adhesion force of individual *Drosophila melanogaster* pupae. The substrate on which the pupa was naturally sticking was fixed on a lab boy using a clamp and brought into contact with a piece of double-sided sticky tape attached to a metal platelet. To detach the pupa from the substrate, a force sensor linked to the platelet by a metal wire was moved up using a motorized micromanipulator. The time-force sensor signal was amplified and converted before being processed in a computer.

328 pupae were measured using our pull-off force measurement set up. In 59 (18%) cases, the pupal case cracked on the dorsal side between the head and the thorax during the pulling process. For 36 (11%) of them, the pupal case covering the ventral part of the head detached from the rest of the pupal case and remained glued to the substrate at the end of the trial, while the complete pupa body was detached together with the posterior part of the pupal case. For the 23 (7%) other cases, the whole animal together with the broken pupal case were detached from the substrate. These 59 (18%) cases were excluded from further analysis. In cases where the animal remained intact, we never observed failure of the pupal case, such as pupal case material remaining attached to the tape or the glue print. Furthermore, two or more trials were sometimes necessary to detach a pupa, when the connection between the double-sided tape and the pupa failed. If the maximal force on the last trial was higher than the previous ones (Fig. 2E), the last trial was used for analysis unless the pupal case was damaged. If the maximal force was higher in one of the first trials, the measurement was excluded, suggesting that the pupa has been partly detached during the first trials (n=23 (7%), Fig. 2F). 11 other cases (3%) were excluded from the analysis: imperfections in the substrates (n=3), pupa not attached ventrally (n=6), pupa not attached, with no glue (n=1) and tape glued to the substrate (n=1). In total, 93 cases (28%) were excluded.

**Figure 2.**
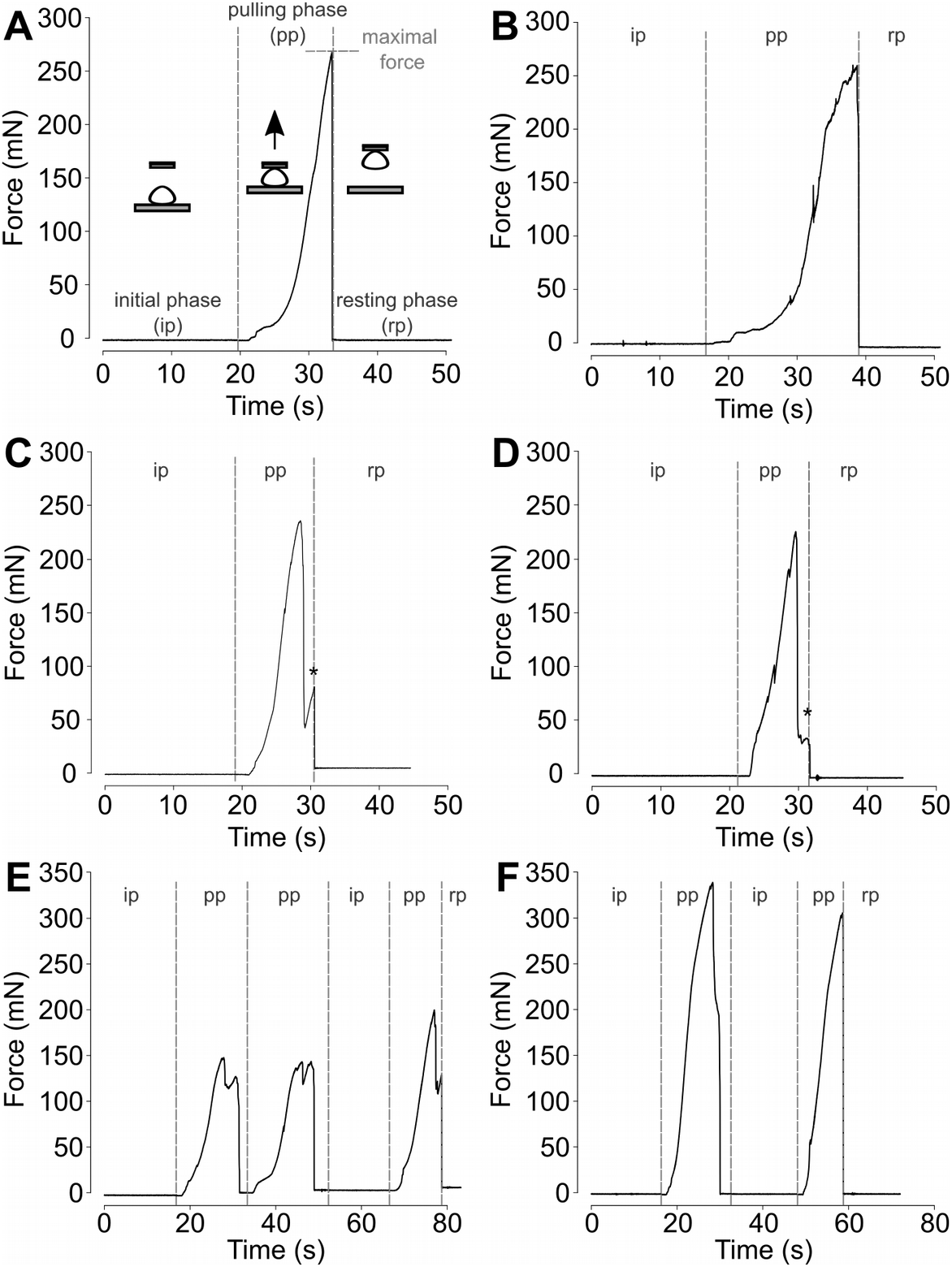
Representative force-time curves obtained in pull-off force measurements. **(A)** Forcetime curve consists of three phases: the initial phase (ip) corresponding to the force before pupa is attached to the tape, the pulling phase (pp) from the time the pupa is attached to the tape to the maximal force, and the resting phase (rp) after detachment. The maximal force corresponds to the force at pupa detachment. Force-time curve corresponding to the attachment of the pupa to the tape (between initial phase and pulling phase) is not shown. Every phase is separated with a vertical dot line. Representative curves of the following cases are shown: **(A)** the tape does not peel from the pupa during the pulling phase and the pupa is detached at once, **(B)** the tape peels from the pupa during the pulling phase and at the end the pupa is detached at once, **(C)** first the posterior part and then the head of the pupa are detaching. Head detachment produces a small peak on the curve (marked with a star), **(D)** the tape peels from the pupa, then the posterior part of the pupa first detaches and then the head of the pupa detaches (marked with a star), **(E)** the tape detaches from the pupa twice and then the pupa detaches from the substrate at the third trial, maximal force is observed during pupa detachment **(F)** tape detaches from pupa once and pupa detaches from the substrate at the second trial, maximal force is observed during tape detachment from the pupa.

We then organized the remaining 235 cases (100%) into four groups. The first group gathers the trials for which the whole pupa detached at once and the tape stayed well fixed to the pupa (n=56 (24%), Fig. 2A). The second group gathers the trials for which the piece of tape started to peel from the pupal case surface during the pulling process and pupa detached at once. For this group, the force-time curve had a negative derivative at some point during the pulling phase (n= 106 (45%), Fig. 2B). The third group gathers the trials for which the whole pupa did not detach at once but the head part stayed attached longer than the rest of the body. In such cases, two peaks were observed on the force-time curve (n=29 (12%), Fig. 2C): the first one corresponds to the main body part detachment and the second one to the head part detachment. The first peak always displayed the maximal force. Finally, the fourth group gathers the trials for which tape started to peel before pupa detachment and for which the whole pupae did not detach at once. For this group, the force-time curve had a negative derivative at some point during the pulling phase and a second peak was observed after the peak corresponding to the maximal force (n=44 (19%), Fig. 2D).

We tested if these four groups having different detachment behavior have an effect on the resulting adhesion force. We performed a two-way ANOVA considering force as the dependent variable. We found that substrates but not groups had a significant impact on pupa adhesion force and that there was no significant interaction between groups and substrates (substrate: F=12.86, P<2e-16, group and interaction group-substrate: P<0.05).

### Microscopy

After pupal detachment, images of glue prints remaining on glass substrates (100 cases) were taken with a Keyence VR 3100 microscope at x40 or x80. Prints areas were measured manually using imageJ (1.50d, java 1.8.0_212, 64-bit) (Schneider et al., 2012). For 1 case, prints were altered before the picture and areas could not be measured. For 11 cases, the red area could not be visualized and was not measured.

One pupa detached from Menzel non-coated glass slide was mounted on aluminum stubs by using double-sided carbon conductive tape (Plano, Wetzlar, Germany). The sample was frozen in liquid nitrogen, sputter-coated with gold–palladium (8 nm thickness) at −140°C, and examined in a cryo-SEM (Hitachi S-4800; Hitachi Ltd., Tokyo, Japan) at −120°C and an accelerating voltage of 3 kV.

To visualize the thickness of the glue at the pupa-substrate interface, *w[*]; P{w[+m*]=Sgs3-GFP}2* larvae were let to pupate as described for measurement assays but on circle cover slips (Thermo Scientific, 32 mm diameter, 0.17 ± 0.01 mm thickness). Untreated (living) samples were directly imaged with Leica SP5 confocal microscope with Plan-Apochromat 40x/1.25 NA objective. Serial optical sections were done every 0.29 μm. Maximal Z projections and Z sections were computed using ImageJ (1.52a, java 1.8.0_112, 64-bit) (Schneider et al., 2012). For the picture of the full attached pupa (Fig. 4A), several images were taken with an Olympus IX83 inverted microscope using a U-Plan FLN 4x/0.13 NA objective and stitched manually.

### Statistical analysis

Statistical differences in pull-off forces between substrates, groups and interactions were tested by two-way ANOVA using R *aov* function (R version 3.4.4, Team, 2013). Statistical differences in pull-force between substrates were tested by one-way ANOVA with the same R *aov* function followed by multiple pairwise comparison tests using Tukey test with R *TukeyHSD* function. Pearson correlation between pull-off force and contact areas were tested using R *lm* function. Effect of humidity, temperature, pressure and age were also tested using R *lm* function.

## RESULTS

### Morphology of the glue

Pupae of *D. melanogaster* attach naturally to substrates on their ventral side via a layer of glue which forms an oval-shaped patch visible on glass slides of approximately 2 mm length and 0.5 mm width (Fig. 3A-B). Glue near the posterior part of the animal usually forms a bigger plug that can spread on the substrate (arrow, Fig. 3B). We performed SEM of pupae detached from non-coated glass slides and found that only thin traces of glue remain on the former contact area (Fig. 3C-G arrow in Fig. 3F). The glue layer covering the pupal case appears to be organized in thin layers (Fig. 3F-G). Air bubbles are observed between the layers and the glue does not fill all the asperities of the cuticle surface (Fig. 3I-J).

**Figure 3.**
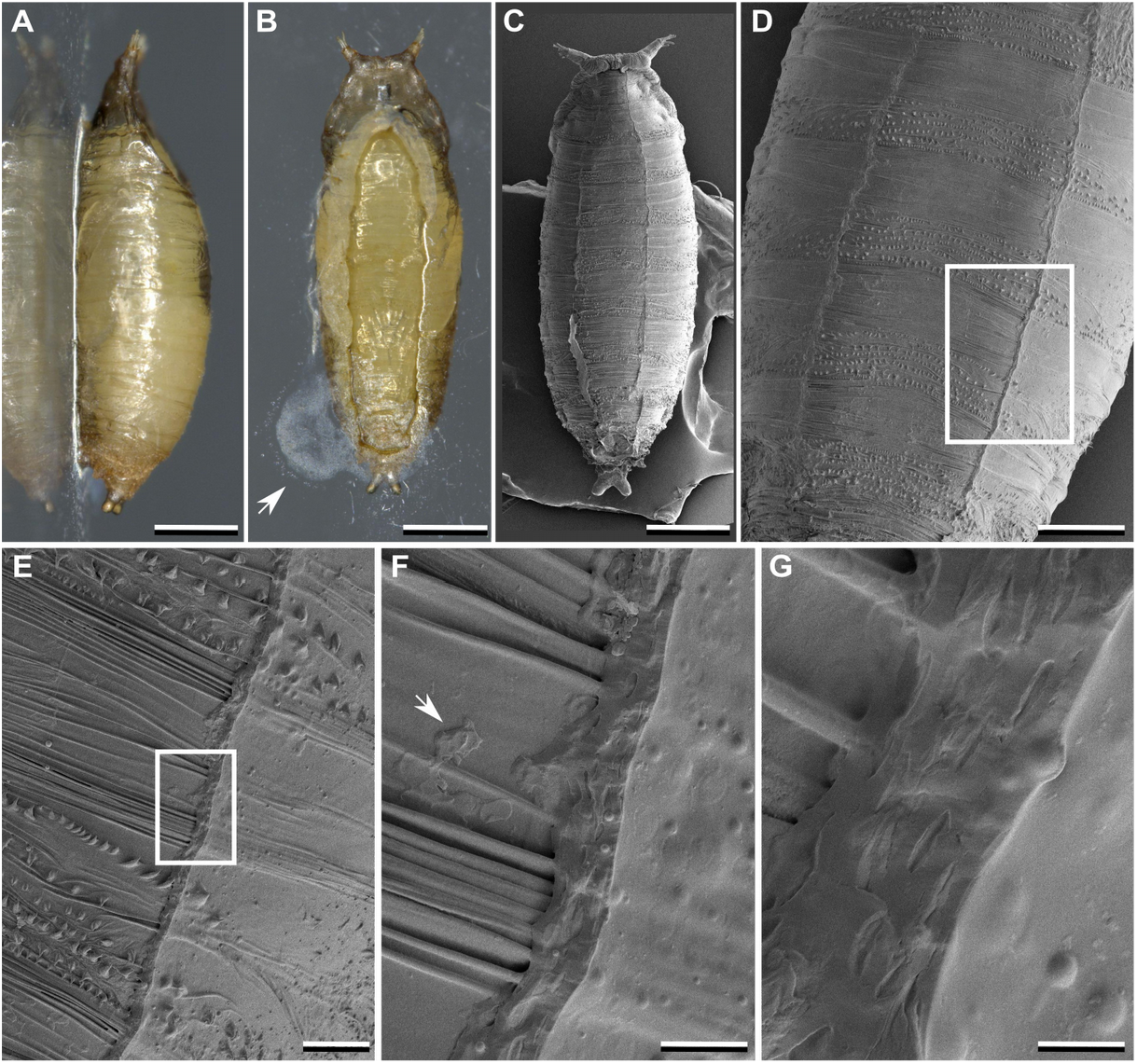
Morphology of *Drosophila melanogaster* glue. **(A,B)** Pupa naturally attached to a glass slide viewed from the side **(A)** and ventrally **(B)**. Glue at the posterior part of the animal sometimes forms a plug on the substrate (**B**, arrow). **(C-G)** Cryo-SEM micrographs of a pupa after detachment from a glass slide. The white squares in **(D)** and **(E)** indicate respectively the location of the images **(E)** and **(F)**. After detachment, glue is not present on the former contact area on the ventral side except thin traces (**F**, arrow) and remains on the sides of the ventral part. Scale bars: **(A-C)** 500 μm, **(D)** 250 μm, **(E)** 50 μm, **(F)** 20 μm. **(G)** 5 μm.

Using confocal microscopy of attached pupae which produce fluorescent glue, we found that the thickness of the glue varies within a single animal from 0 to about 20 μm due to the irregularities of the pupal case and its barrel shape (Fig. 4). We note also that this fluorescent glue could have slightly different properties than wild-type glue.

**Figure 4.**
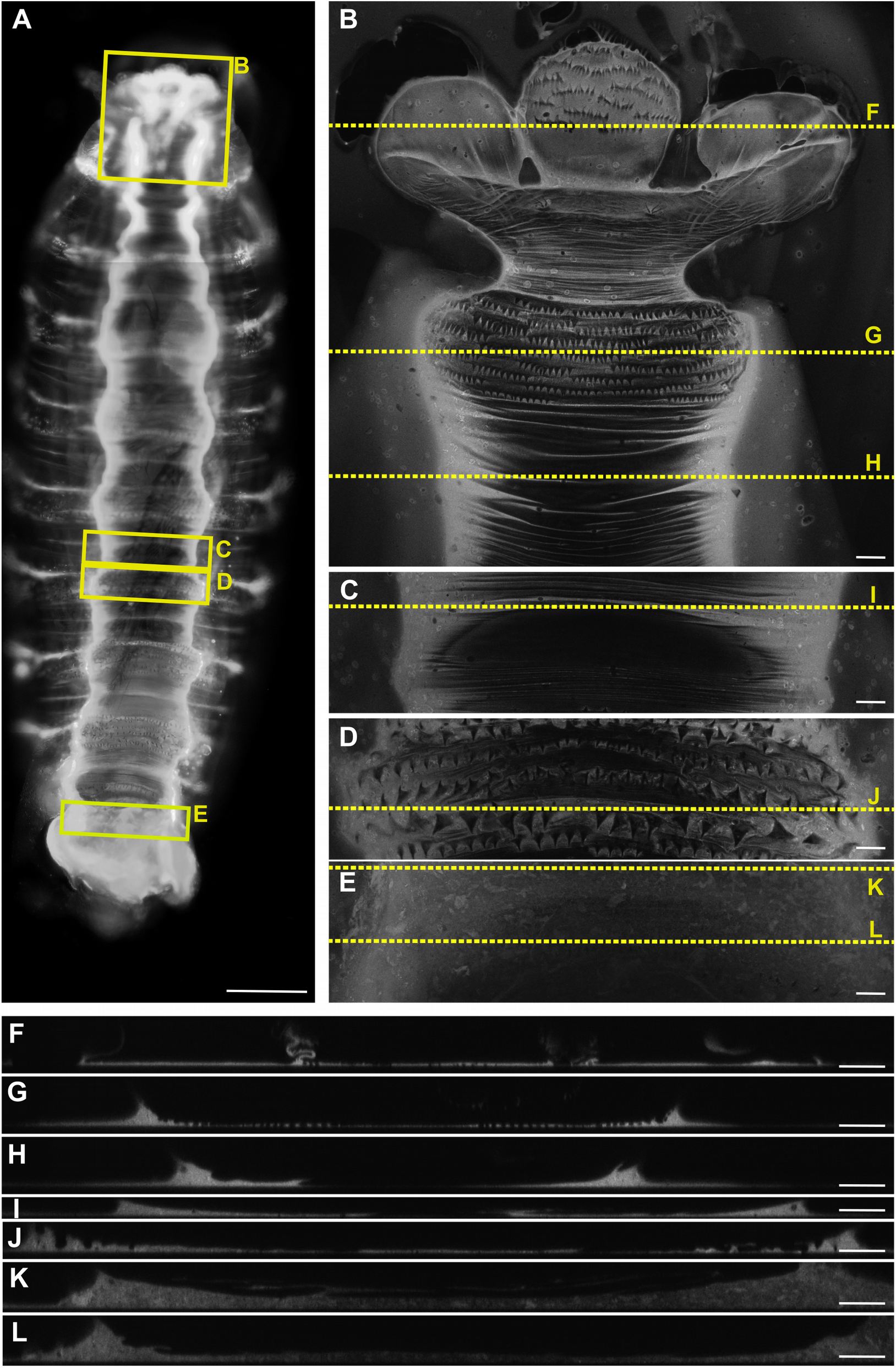
Observation of the fluorescent solidified glue produced by a *Drosophila melanogaster* pupa *w[*]; P{w[+m*]=Sgs3-GFP}2*. **(A)** Fluorescent microscopy of the attached pupa viewed from the ventral side. **(B-E)** Confocal microscopy of anterior **(B)**, middle **(C,D)** and posterior **(E)** ventral regions of the attached pupa. Maximal Z projections are shown for each region. **(F-L)** Z sections of the pupa-substrate interface. The yellow squares in **(A)** indicate the locations of images **(B-E)**. The dashed red lines in **(B-E)** indicate the Z sections projection axes. **(A-E)** Anterior is up. **(F-L)** Substrate is at the bottom. Scale bars: **(A)** 200 μm, **(B-E)** 20 μm, **(F-L) 20** μm.

### Pupal adhesion on different substrates

We performed pull-off force measurements using pupae naturally attached to 11 different substrates: non-coated glass (from Roth and from Menzel), PLL-coated glass, PLL-PEG-coated glass, oxygen-activated glass, Teflon, and Spurr epoxy resin with 5 different roughnesses. We measured the contact angle of these 11 substrates (Table S2). Contact angle values ranged from 11° (highly hydrophilic substrate) to 112° (highly hydrophobic substrate). In total, 328 pupae were measured. We excluded 93 cases for which adhesion measures were not reliable, for instance when the pupal case was damaged during the assay or when several trials with the same pupa disrupted its adhesion (see Methods).

Medians of the pull-off forces on glass-substrates (non-coated, PLL-coated, PLL-PEG-coated and oxygen-activated glass) ranged from 184 mN (oxygen-activated glass, SD= 78) to 229 mN (PLL-PEG-coated glass, SD= 122) while for Teflon, force was 42 mN (SD= 20) (Fig. 5). Only pull-off force on Teflon was significantly different from forces obtained on other substrates (oneway ANOVA F=12.92, P<2e-16, followed by all pairwise comparison Tukey-Test, P<0.001 for all comparisons with Teflon). Similarly, pull-off force medians on resin with different roughnesses ranged from 151 mN (P80, SD= 82) to 269 mN (1MIC, SD= 105) and no statistical differences were found (from same ANOVA and Tukey tests). During our assays, the ranges for humidity, temperature, pressure and age of pupa were the following: 34.4 – 58.8 %, 23.5 – 27.9 °C, 1005 – 1026 mb, 3.5 – 23 h after deposition of the wandering L3 larva on the substrate. No effect of humidity, temperature, pressure and age of pupa was found (multiple linear regression with force as dependent variable and substrate, humidity, pressure, age as independent variables).

**Figure 5.**
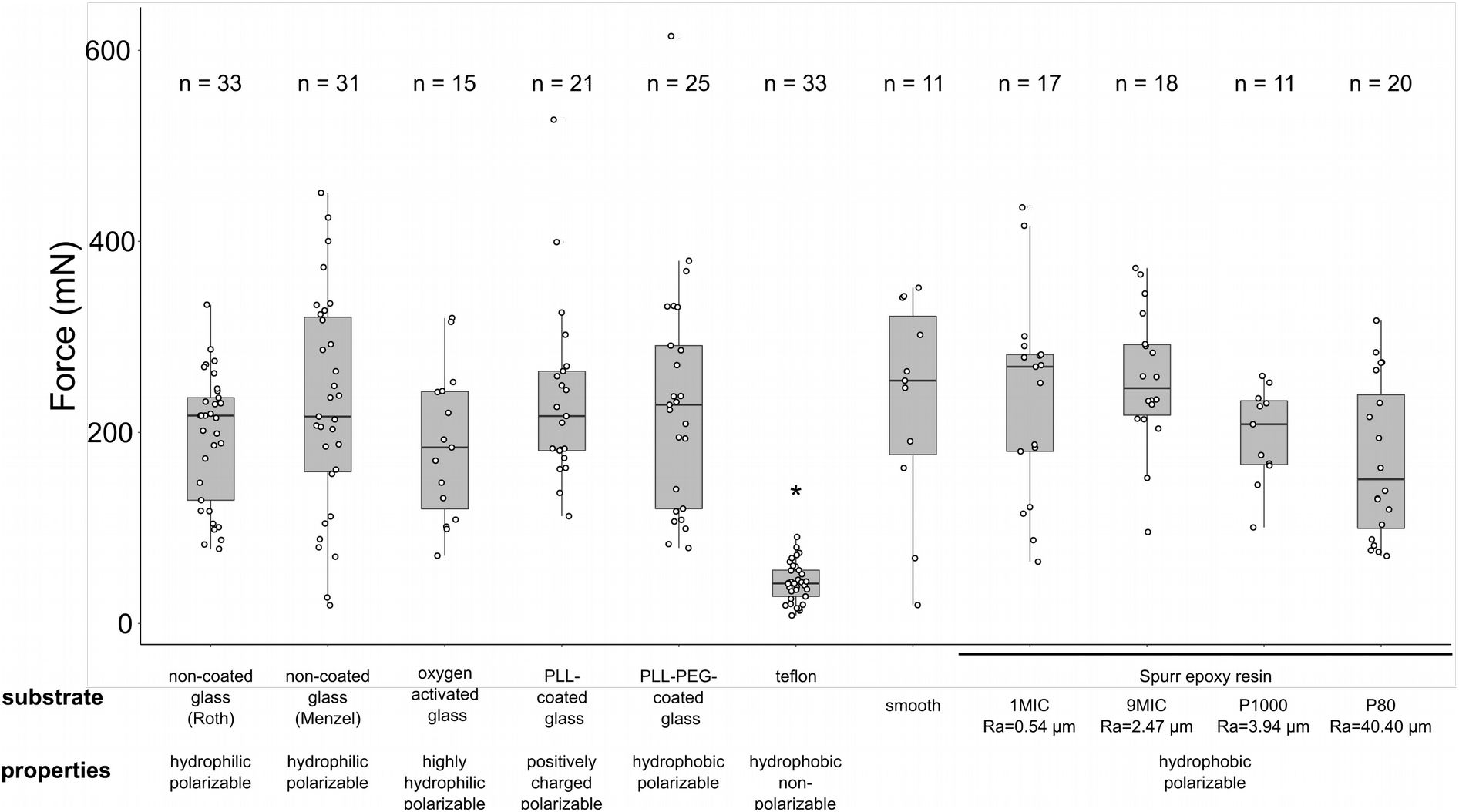
Force required to pull-off pupae from different substrates. Each dot corresponds to a single pupa and n indicates the total number of pupae tested for each surface. Ends of the boxes define the first and third quartiles. The black horizontal line represents the median. The vertical line on the top of the box extends to the largest value no further than 1.5 * IQR from the upper hinge of the box. The vertical line on the bottom of the box extends to the smallest value at most 1.5 * IQR of the hinge. (IQR: inter-quartile range is the distance between the first and the third quartiles). Data beyond the end of these lines are “outlying” points. Asterisks indicate significant difference (P<0.001) between Teflon and all the other substrates. Ra values correspond to roughness averages for the different resin substrates.

In conclusion, we found that the pull-off force is independent of substrate surface chemistry (except Teflon) and of roughness in wide range (80 nm – 40.40 μm).

### The glue-substrate interface

After pull-off force measurements, glue prints were analyzed for the five glass-type substrates. For these substrates, three types of prints can be defined: (1) the glue fully remained on the substrate (Fig. 6A), (2) the glue partly detached from the substrate (Fig. 6B) or (3) the glue went off completely with the pupa (Fig. 6C). Most of the time, the glue was partly detached from the substrate (n= 95/124). For the five glass-type substrates, print types (but not substrate types) have a significant impact on pupa adhesion (two-way ANOVA, print types: F=5.6, P=0.0013). When the glue completely went off with the pupa, corresponding pull-off force was significantly lower than force obtained when glue was partly detached (134 mN, SD= 62 compared to 231 mN, SD= 102, Tukey test P<0.05) (Fig. 6D). On Teflon, the glue always went off with the pupa (n=33/33).

**Figure 6.**
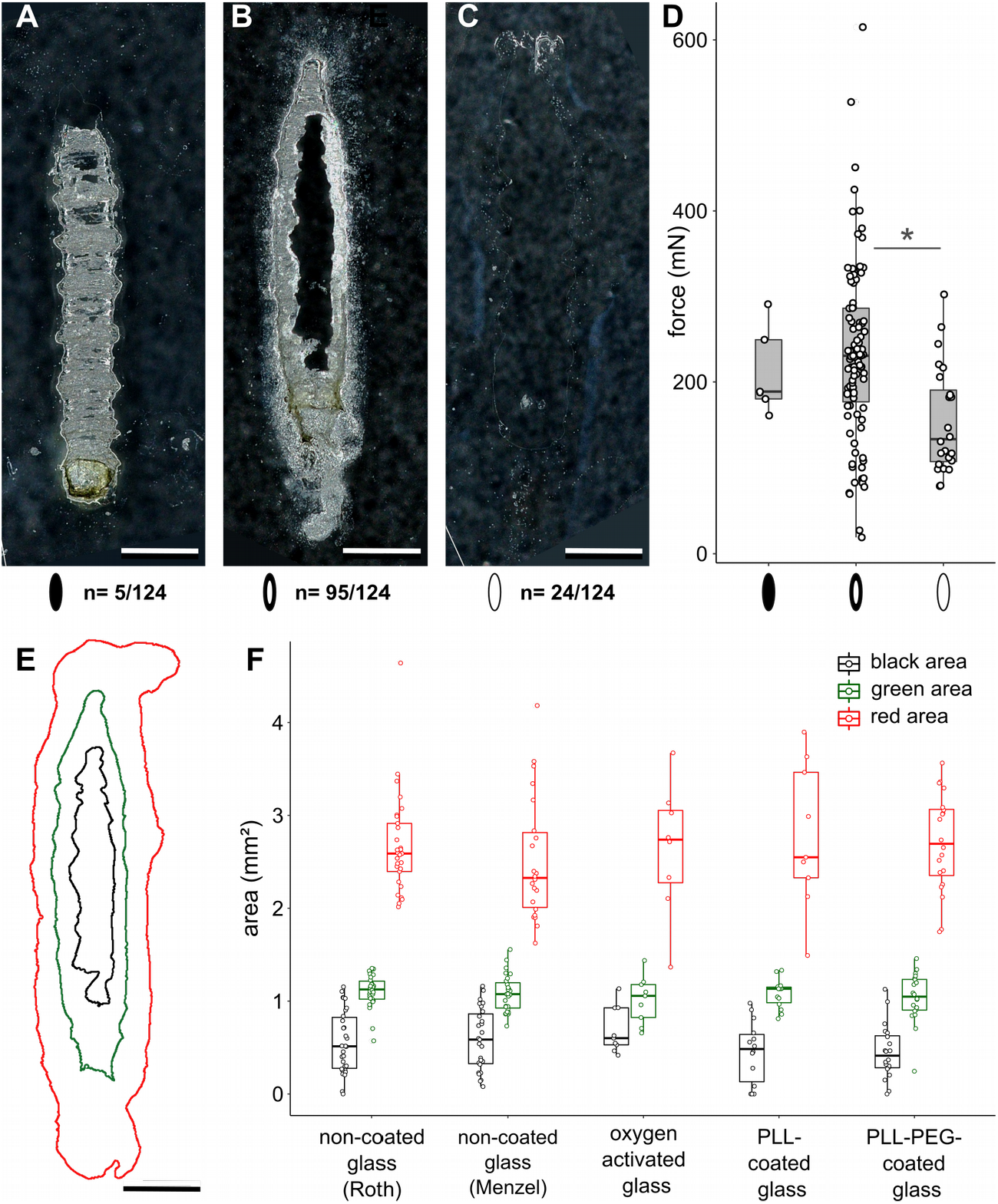
Glue-substrate interface on glass-type substrates. **(A-C)** Examples of typical glue prints obtained after pull-off force assays on glass-type substrates: **(A,C)** PLL-coated glass, **(B)** non-coated glass (Roth). Three cases are defined: **(A)** when the glue fully remained on the substrate, represented as a black oval, **(B)** when the glue partly detached from the substrate, represented as a white oval with thick black contour, **(C)** when the glue completely went off from the substrate, represented as a white oval. n indicates the total number of glue prints obtained for each case. **(D)** Box plots representing the force required to detach pupae depending on the print types defined in **(A-C)**. One dot corresponds to a single pupa. Asterisk indicates significant difference (P<0.05) between cases where the glue partly detached from the substrate and cases where the glue completely went off from the substrate. **(E)** Scheme of the various glue prints with the glue-pupa interface delimited in green, the glue-glass interface delimited in red and the border of the glue removed from the glass after pull-off adhesion test in black. **(F)** Box plots showing the size of the three areas defined in (E) for the five glass-substrates. Only prints where the glue fully remained and partly detached from the substrates were analyzed (see Methods). Scale bars: **(A-C,E)** 500 μm. Box plots are defined as previously (Fig. 5).

Furthermore, from the glue print three distinct areas can be defined. The pupa-substrate interface area corresponds to the area where the pupa was in contact with the substrate (green outline in Fig. 6E). The glue-substrate interface area (red outline in Fig. 6E) corresponds to the total surface of the substrate covered by the glue. After pupa detachment, a third area where glue went away with the pupa can be defined (black outline in Fig. 6E). For all cases (as in Fig. 6A,B) where the glue did not go off completely with the pupa, the three areas were measured from the print images of the five glass-type substrates. A relatively high variation in glue-substrate interface area was observed (from 1.3 to 4.6 mm^2^) compared to the pupa-substrate area which was relatively invariant, around 0.2 to 1.5 mm^2^ (Fig. 6F). No difference was found in areas between substrates (one-way ANOVA for each area, P>0.05) (Fig. 6F). No correlation was found between pull-off force and any of the three areas when combining the five substrates or when testing each substrate individually (linear regression for each area, P>0.05) (Fig. S1).

## DISCUSSION

*Drosophila* glue spreads over an external substrate and sticks the pupa’s ventral part to the substrate, forming an oval-shaped interface (Fig. 3B). From SEM images, we observed flat holes that seem to correspond to air bubbles that were trapped between different layers of glue. Several layers may have been deposited successively on the surface of the pupa by the back and forth movements of the larvae during expectoration (Fraenkel and Brookes, 1953). The presence of bubbles and the fact that the glue does not fill all the asperities of the cuticles suggest that the glue is relatively viscous when it comes out. It is known that the glue is produced by the salivary glands and is secreted by the mouth (Fraenkel and Brookes, 1953). However, on the glue prints we sometimes observed a plug around the posterior tip of the larva (Fig. 3B). Recently, Benova-Liszekova et al. (2019) reported that a few larvae did not empty their guts entirely before pupariation. One possibility is that this plug might be a posterior secretion from the digestive tube, which mixes with the anteriorly secreted glue.

During our pull-off force assays, the piece of tape used to stick the pupa in order to pull it peeled out from the pupal case in 153 of the 235 analysed cases, meaning that the adhesion strength between the double-sided tape and the pupal case is close to the adhesion strength between the glue and the pupa. In our experiments, the tape thus adheres to the pupal case just enough for pupa glue measurements. Furthermore, in 76/235 cases, the posterior part of the animal first detached from the substrate before the head detached. In these cases, the pull-off force was probably not applied exactly at the center of the pupa but more posteriorly due to the asymmetrical shape of the pupa. Furthermore, the pupal case is more fragile around the puparial opercular seam, a seam which runs across the front of the puparium and extends along the sides, and which will break when the adult will hatch (Tyler, 1994). In addition, when using forceps to extract the animal from its pupal case, the pupal case generally splits into annular strips (Held, 1992). The zone of breakage that we observed on the pupal case in our pull-off assays therefore corresponds to the most fragile region of the pupal case.

In any case, no significant difference was found between cases for which pupae detached at once and cases where the posterior part detached before the head part. This suggests that stronger head adhesion does not affect the maximal pull-off force.

Pull-off forces required to detach pupae from substrates other than Teflon ranged from 151 mN (P80, SD=20) to 269 mN (1MIC, SD=105). Using pupa-substrate interface (green outline in Fig. 6E) as a measure of contact area between the glue and the substrate, we found a median contact area of 1.10 mm^2^, which leads to an adhesive strength ranging from 137 to 244 kPa. Few similar studies have been carried on insect egg adhesive strength. Two of them reported higher values (1-2 MPa for *Opodiphthera* eggs on wood (Li et al., 2008), 4.3-12.2 MPa for whitefly eggs on rose leaves (Voigt et al., 2019), one study found equivalent values (38.8-271.3 kPa for beetle *C. asparagi* on plant surface (Voigt and Gorb, 2010)) or another lower adhesive strength (13.9-97.8 kPa for the moth *C. pomonella* eggs on fruits (Al Bitar et al., 2014)). To our knowledge, our study is the first one to report adhesion strength measurements for Diptera pupa.

No statistical differences in pull-off forces were found between the different substrates except for Teflon (median pull-off force is 42 mN, SD= 20), suggesting that the glue can stick to hydrophobic substrates as well as hydrophilic substrates. We can hypothesize that proteins of the glue are polarizable, which allows the glue to stick strongly to polarizable substrates that are differently charged and not to Teflon, which is not polarizable. Indeed, Sgs proteins are highly charged. They are mostly composed of positively charged amino acids and they are highly glycosylated with negatively charged sugars (Beckendorf and Kafatos, 1976; Korge, 1977).

No differences were found on resin substrates with different roughnesses. We expected that pull-off force would increase with roughness as contact area on rough substrate is bigger. Furthermore, we did not find correlation between force and contact area for any of the substrates used. Both observations can be explained by a possible critical crack initiation stress between the pupal case and the glue. Indeed, we observed on the glue prints on glass-type substrates that the glue from the pupa-substrate contact area is partly detached, suggesting that rupture occurs between the glue and the pupal case. A similar phenomenon may occur on resin substrates, whereas on Teflon the glue completely goes off from the substrate, suggesting that the rupture occurs between the glue and the Teflon. Thus, in all the tested substrates except Teflon, the bond between the glue and the different substrates appears to be stronger than the bond between the glue and the pupal case. It is thus possible that our adhesion tests measure the adhesive force of the bond between the glue and the pupal case, which would explain why we observed the same adhesion strength for all substrates except Teflon. As an alternative explanation for the same adhesiveness measured on all tested substrates except Teflon, it is possible that the animal modulates the amount of glue it produces, or the way it spreads the glue over the substrate and its body. It would be interesting to do adhesion measurement on isolated glue. However, it is challenging to collect the glue as it is secreted in very small volume and it polymerizes within 3-5 min. after expectoration (Beňová-Liszeková et al., 2019).

Pull-off force values were rather variable between trials for a given substrate (for example from 19 to 451 mN on Menzel non-coated glass). Such a wide range of values could be due to individual variability. Pupae weight could not be deduced accurately by subtracting the final force from the initial force because pupal weight was within the noise of force values. However, individuals from the same population as the pupae used for trials were weighted instead. We found that pupal weight averaged 1.4 ± 0.3 (SE) mg (n=24). The weight variation is negligible compared to the force variation observed within substrates. Surprisingly, considering this average weight and an average force of 217 mN (mean of all trials kept for analysis except trials on Teflon), the glue holds about 15 500 times the weight of a pupa.

## Conclusions

For the first time, we report here measurements of *Drosophila* pupal adhesion strength. We present a pull-off force test to measure pupa adhesion which could be used in the future to explore pupa adhesion in various strains and species of flies. With the powerful genetic tools available in *Drosophila melanogaster*, we now plan to assess the adhesive function of the glue genes. The use of *D. melanogaster* as a model organism for the study of bioadhesives is very promising as it makes it possible and easy to manipulate the composition of the glue.

## Supporting information

Borne_2020_table

## Abbreviations

PLL: Poly-L-Lysine
PLL-PEG: Poly-L-Lysine-Polyethyl glycol
SEM: Scanning Electron Microscopy

## Acknowledgements

We acknowledgeT. Roeder and C. Sandberg (Kiel University, Germany) for help with fly culture, S. De Beco (Université de Paris, Paris) for help with PLL-PEG coating and L. Corté (Mines-ParisTech, ESPCI, Paris) for discussions. We also thank the ImagoSeine facility, member of the France BioImaging infrastructure supported by the French National Research Agency (ANR-10-INSB-04, «Investments for the future» and N. Moisan for technical assistance with confocal microscopy. Stocks obtained from the Bloomington Drosophila Stock Center (NIH P40OD018537) were used in this study.

## Supplemental data of Borne et al., 2020

**Table S1. raw dataset: Borne_2020_glue_table_S1.csv**

**Table S2.** Contact angle measurements (°) for the different substrates. Contact angles were measured for each substrate. All the measurements were done before the pull-off force measurements except for PLL-PEG-coated glass.

| Substrate | mean contact angle (°C) | standard error |
| --- | --- | --- |
| 1-MIC resin | 112.68 | 0.47 |
| 9-MIC resin | 78.63 | 1.89 |
| P1000 resin | 109.19 | 0.89 |
| P80 resin | 103.27 | 1.29 |
| smooth resin | 84.11 | 1.93 |
| PLL-coated glass | 55.15 | 1.28 |
| PLL-PEG-coated glass | 38.65 | 1.06 |
| oxygen-activated glass | 37.03 | 1.39 |
| non-coated glass (Menzel) | 28.86 | 0.91 |
| non-coated glass (Roth) unwashed | 20.05 | 0.57 |
| non-coated glass (Roth) washed | 11.84 | 1.40 |
| teflon | 96.64 | 1.88 |

**Figure S1.**
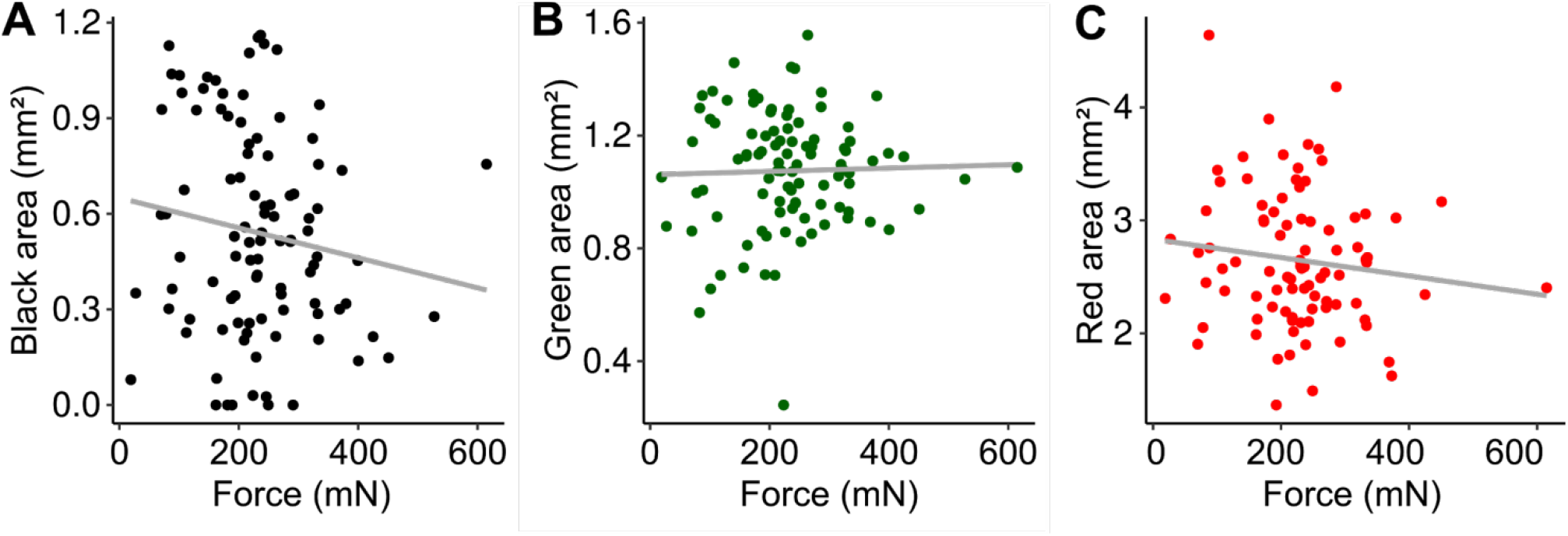
Force in function of (A) black area, (B) green area and (C) red area. Each dot represents one print on glass-type substrates. The grey lines represent the linear regressions: **(A)** r^2^= 0.01168, y= 257.69 – 46.03x, p=0.144; **(B)** r^2^= −0.009391, y= 218.05 + 13.73x, p=0.7793; **(C)** r^2^= 0.004791, y= 279.36 – 19.97x, p= 0.2369; where C is the adjusted C.

